# Human infection with an ape *Plasmodium vivax*-like parasite

**DOI:** 10.64898/2026.10.23.757543

**Authors:** Tue Sparholt Jørgensen, Paul M. Sharp, Jonathan Szeps, John White, Thomas James Booth, Henrik Vedel Nielsen, Christen Rune Stensvold

## Abstract

An unusually divergent *Plasmodium vivax* parasite was reported in a Swedish patient with recent travel. Using whole-genome sequencing, we show that the infection was caused by a strain of ape *P. vivax*-like, providing evidence of the zoonotic potential of these parasites that infect wild gorillas and chimpanzees in central Africa.

**Summary line:** Whole-genome sequencing identified a naturally acquired human infection with a malaria-causing ape *Plasmodium vivax*-like parasite, providing further evidence for the zoonotic potential of these parasites.

---

While relatively rare in central Africa, *Plasmodium vivax* is the most widespread human malaria parasite outside Africa. Over the past 16 years, DNA sequences of *P. vivax*-like parasites have been isolated from samples from wild individuals of all four species of African apes; *i*.*e*., western and eastern gorillas, chimpanzees and bonobos (1-4). These ape parasites harbor about 10 times as much genetic diversity as the human *P. vivax* parasites, suggesting that the human parasites have undergone a genetic bottleneck (2, 3, 5). It is unclear whether the ape and human strains of *P. vivax* have become separate species, and the ease by which ape strains can infect humans remains unknown. Indeed, only a single case of human infection with ape *P. vivax-*like has been reported to date (6). That report was based on mitochondrial DNA and short fragments of 2 nuclear genes, limiting robust phylogenetic placement. We recently reported a diagnostically challenging malaria case from Sweden in a patient with an extensive travel history, in whom amplicon-based next-generation sequencing (NGS) was required to identify the *Plasmodium* species responsible (7). Partial A-, S-, and O-type SSU rRNA sequences were obtained, each of which clustered phylogenetically with reference sequences from *P. vivax*. However, these sequences were somewhat divergent from those of human *P. vivax* strains. Therefore, we moved on to fully characterizing this strain by determining the complete genome sequence. Phylogenetic analyses reveal that the strain lies within the radiation of ape *P. vivax*-like strains, quite distinct from human *P. vivax*, providing genomic evidence of zoonotic transmission.

## The Study

### Technical details are provided in the Appendix

Only 0.001% of the original patient DNA sample ‘Pvl-KarSSI’ (called ‘52118’ in the previous paper, (7)) was classified as *Plasmodium* derived, which would normally prevent further analysis. We used selective whole-genome amplification (SWGA) (8) to enrich the *Plasmodium* genome ∼6,000-fold over the background of human DNA, enabling cost-efficient DNA sequencing. Illumina and Oxford Nanopore data were generated from the SWGA sample and used for de novo hybrid genome assembly (9). Publicly available *P. vivax* assemblies (n=16) and ape *P. vivax*-like genomic datasets (n=18) were downloaded from GenBank and the Sequence Read Archive at NCBI, and filtered for co-infections (10). Shared single-copy orthologous protein sequences were used for genome-wide comparisons of all genomes (11).

Assembly of Pvl-KarSSI generated a genome of 21.1 Mbp in 473 contigs with an N50 of 111 kbp. This corresponds to 93% of the chromosome length of the human *P. vivax* reference Sal1 (GCF_000002415.2). We identified 93% of 4203 complete core *Plasmodium* protein coding genes in Pvl-KarSSI, similar to the three available *P. vivax*-like reference assemblies Pvl01 (87% complete), PvSY43 (92% complete) and PvSY56 (93% complete). Based on the size and gene content, we conclude that our assembly is of similar quality to the three references.

Phylogenetic analyses based on 356 concatenated conserved BUSCO proteins (12) place the Pvl-KarSSI sample among the ape *P. vivax*-like strains rather than with human *P. vivax* (Figure 1). The tree has three major clades, one comprised of globally distributed human *P. vivax* strains including samples from Asia, Africa, and the Americas (see Supplementary Table 1 and the Appendix), and two comprised of ape *P. vivax*-like strains, organized as previously described in group 1 and group 2 (5). All three main clades have a support of 1, indicating a solid placement of all sequences. Phylogenetic trees estimated from reduced protein datasets present in 100% of samples, and excluding samples with low completeness, showed identical main clades (see Appendix). Pvl-KarSSI is contained in the ape *P. vivax* ‘group 2’ clade, clustered significantly with a strain from a gorilla sample from Gabon (Figure 1). There does not seem to be any clear host species split between groups 1 and 2, since both groups have been found in samples from both chimpanzees and gorillas. The limited sequence data from the previous human sample of ape *P. vivax*-like, the “traveler CAR” sample from 2013 (6), are not sufficient to determine if they stem from the same lineage as Pvl-KarSSI. In the short overlapping genomic windows, the TravelerCAR and Pvl-KarSSI samples are not identical (see Appendix).

**Figure 1.**
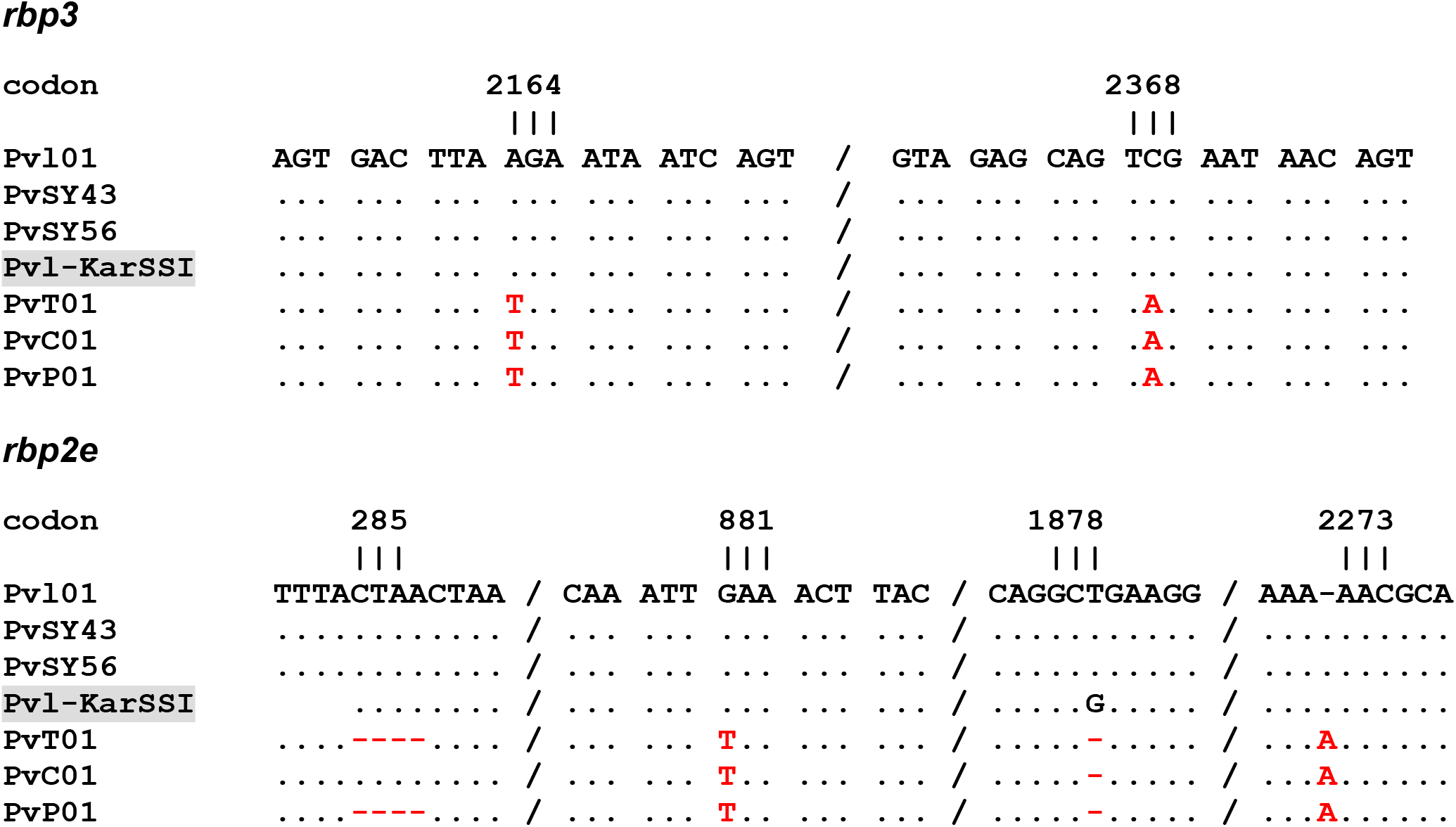
Unrooted phylogenetic tree estimated from 356 concatenated core BUSCO proteins with at least 90% presence, listed in the Appendix. The approximately-maximum likelihood, using FastTree in getphylo, was applied with the JTT+CAT model of protein evolution. Only SH-like support values 0.98 or higher are displayed. Colored squares denote the host from which the sample was obtained. The new strain described here, Pvl-KarSSI, is in bold.

*P. vivax* genomes include ten members of the *rbp* gene family, encoding reticulocyte-binding proteins involved in red blood cell invasion. In all strains of *P. vivax* from humans, the three genes *rbp2d, rbp2e* and *rbp3* are pseudogenes, whereas strains of *P. vivax*-like from apes lack the inactivating mutations (3). We searched for these genes in the Pvl-KarSSI genome sequence. The *rbp2d* gene was not present in the assembly. In *P. vivax*, the *rbp2e* gene comprises two exons, of 55 bp and 8144 bp, respectively. In our assembly, only the last 7347 bp of exon 2 are found, at the beginning of a contig. Within this region, most strains of *P. vivax* from humans carry 2-4 indels leading to frameshifts, as well as a premature stop codon (3); none of these inactivating mutations are present in Pvl-KarSSI (Figure 2). A complete sequence of *rbp3* is present in our assembly. More than 80% of strains of *P. vivax* from humans have two premature stop codons in *rbp3* (3), but again neither mutation is found in Pvl-KarSSI (Figure 2). In conclusion, the sequences of *rbp2e* and *rbp3* in Pvl-KarSSI independently support assignment of Pvl-KarSSI to the ape *P. vivax*-like lineage.

**Figure 2.**
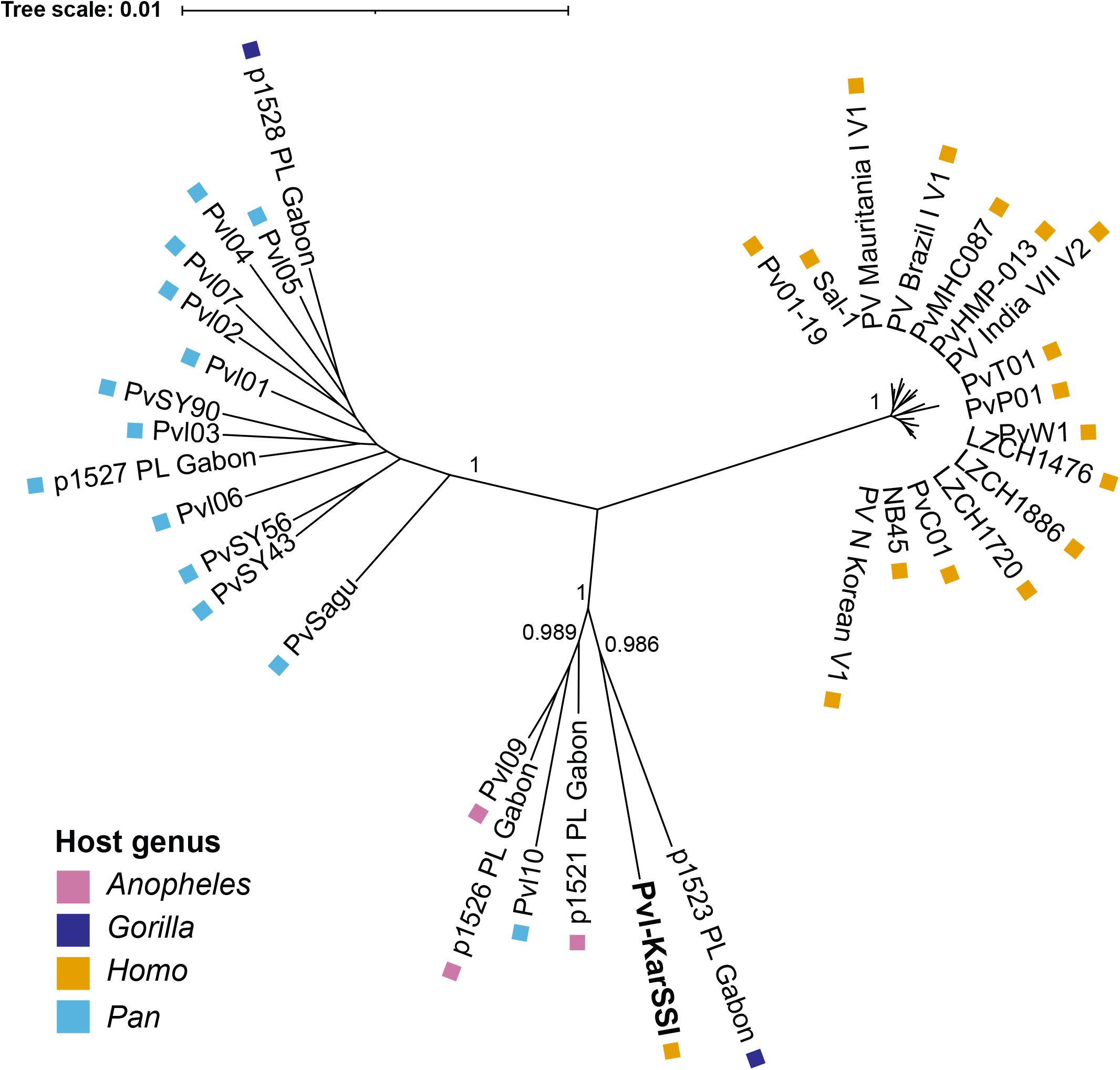
Sites of mutations (in red) that make *rbp3* and *rbp2e* pseudogenes in strains of *P. vivax* from humans are not mutated in Pvl-KarSSI, the strain characterized here. Pvl-KarSSI is compared to three strains of *P. vivax*-like from chimpanzees (Pvl01, PvSY43 and PvSy56) and three strains of *P. vivax* from humans (PvT01 from Thailand, PvC01 from China, and PvP01 from Indonesia). In *rbp3*, codons 2164 and 2368 encode Arg and Ser, respectively, but are premature stop codons in *P. vivax* from humans (at least one of these mutations was found in around 97% of 227 strains analyzed by Loy et al. 2018). In *rbp2e*, codon 881 encodes Glu, but is a premature stop codon in *P. vivax* from humans; additionally, deletions around codons 285 and 1878, and an insertion in codon 2273, all shift the reading frame (at least one of these mutations was found in around 93% of 162 strains analyzed by Loy et al. 2018). In Pvl-KarSSI, the mutation at the third position of codon 1878 of rbp2e is synonymous. The *rbp2e* sequence from Pvl-KarSSI is missing codons 1-284 at the end of the assembled contig.

## Conclusions

Whole-genome sequencing documented that the parasite infecting this patient belongs to the ape *P. vivax*-like group 2. The only previous report of human infection with ape *P. vivax*-like parasites, from 2013, analyzed only mitochondrial and limited nuclear DNA fragments, preventing whole genome-based comparisons. Thus, the case presented here is the first comprehensively documented naturally acquired human infection with ape *P. vivax*-like malaria parasite.

This case highlights the value of two facets of sequencing-based analysis for understanding parasite infection. First, the amplicon-based NGS results from the sample presented previously were fundamental in identifying Pvl-KarSSI as particularly interesting. And second, the technological advance which SWGA represents enabled us to generate sufficient coverage of the ape *P. vivax*-like genome to comprehensively place this sample within the ape *P. vivax*-like diversity.

The patient reported travel involving field work in wildlife habitats in the Republic of the Congo, in Rwanda, and in Indonesia at dates 9 months, 8 months, and 4 months before the diagnosis in 2025. The only visit to a known ape habitat was a November 2024 trip to rainforest near Bomassa, in the Republic of the Congo, near a site where both chimpanzees and gorillas have previously been found to be infected (2). This represents a plausible period of exposure, because human *P. vivax* parasites can display a long incubation phenotype, with documented cases of 8-9 months between infection and symptoms (13).

The genomic data placing this parasite within the ape *P. vivax*-like radiation provides clear evidence that those parasites have zoonotic potential. The finding raises questions about the frequency of such spillover infections and the extent to which they may remain unrecognized by routine malaria diagnostics.

## DATA AVAILABILITY

Data are deposited under study accession PRJNA1507507. The genome assembly of Pvl-KarSSI is available under accession JCEGHI01. *P. vivax*-like assemblies and annotation files are available via DOI10.6084/m9.figshare.33241578

## ACKNOWLEDGEMENTS

We thank Katja Wyss, Silvia Botero Kleiven, Kristoffer Kiil, and Virginie Rougeron for inspiring discussions on the sample, sequence analysis and phylogenetic placement, and Martin Schou Pedersen, Anna Rønberg Schmidt-Nielsen, and the SSI core sequencing facility for excellent technical support. This work was supported by the Novo Nordisk Foundation through the grants NNF22OC0078997 and NNF23400085294. Finally, we want to thank the patient for giving written informed consent to this publication.

