## Appendix for "Human infection with an ape *Plasmodium vivax*-like parasite"

**Appendix for the manuscript Human infection with an ape *Plasmodium vivax*-like parasite**

Summary: Here we present the first genome from a human case of ape *Plasmodium vivax*-like, from a European traveler. We document the placement as an ape *P. vivax*-like using core protein trees and analysis of the genes encoding two reticulocyte binding proteins. This document details methods used for the placement, as well as supplementary results which establish the sample Pvl-KarSSI as a single infection without evidence of contamination, recombination, or other issues, and which support the validity of the main phylogenetic tree using reduced datasets and individual protein phylogenetic trees built from the samples in Supplementary Table 1. Further, details on the generation of assemblies of previously published samples of ape *P. vivax*-like is described.

**Content:**

Supplementary Information	Detailed methods and results

Supplementary Figure A	Example phylogenetic trees of individual proteins

Supplementary Figure B	Reduced phylogenetic tree from 100% presence, only HQ samples

Supplementary Figure C	Comparison between Pvl-KarSSI and TravelerCAR

Supplementary Table 1	*P. vivax* and *P. vivax*-like samples used

Supplementary Table 2	Primer list (Pvset1920)

Supplementary Table 3	List of taxIDs used for krakenuniq read filtering

Supplementary Table 4	Name and description of 356 BUSCOs used for Figure 1

Available at Figshare reviewer link https://figshare.com/s/9b0c4ea2571e2ac0f97d:

Supplementary Dataset I	Annotated assemblies in GenBank file format

Supplementary Dataset II	Mapping bam file used for polymorphism determination

Supplementary Dataset III	Getphylo results

Supplementary Dataset IV	Consensus nucleotide fasta of the mitochondrion and 18S rDNA fragments from Pvl-KarSSI used for Supplementary Figure C

**Supplementary Information**

Detailed methods

After initial DNA QC using tapestation (genomic DNA screentape), qubit (dsDNA HS), and nanodrop on the original, very thin, 0.5ng/ul EDTA blood extraction, we generated 2 x 150 nt Illumina Nextseq1000 data from a Hackflex (1) library, yielding 1.3M paired end reads. Of these, 16, or 0.001% were classified as *Plasmodium*. Selective whole genome amplification (SWGA) was then performed according to Cowell et al., 2017 (2). Briefly, 3.5uM primer set pv1920 (TAG Copenhagen, Supplementary Table 2), 30U phi29 polymerase (NEB M0269S), 1x buffer, 4mM dNTPs. 4% bovine serum albumin (0.08 % w/v) was used instead of the 1% used by Cowell et al. The total reaction volume was 50 ul. The SWGA reaction was incubated at gradually reduced temperature in a ramp down from 35°C to 30°C (10 min per degree), followed by 16 h at 30°C and 10 min at 65°C for inactivation. Another Illumina Hackflex library was generated directly from the reaction mix, and sequenced similarly to above. This yielded 6.7M *Plasmodium* illumina reads out of 91M total reads (7.3%), the equivalent of a ca 6000x amplification over the human background. SWGA DNA was debranched using the NEB T7 Endonuclease I according to manufacturer instructions. Debranched DNA was used to construct an Oxford Nanopore ligation library (SQK-LSK114-96), which was sequenced on an Oxford Oxford Nanopore MinION 10.4.1 flowcell (dna_r10.4.1_e8.2_400bps_sup@v5.2.0). KrakenUniq (v. 1.0.4) (3) with the microbialDB of bacteria, parasites and human was used for filtering out non-*Plasmodium* illumina reads using the TAXIDs 5820 and 0 (Plasmodium and unclassified). For Oxford Nanopore data, non-*Plasmodium* was filtered out by first mapping on both the *P. vivax* reference SALI (4) and the human genome (GRCh38.p14), and then using Samtools (5) to keep only reads which had best map on the *Plasmodium vivax* reference genome SalI (GCA_000002415.2_ASM241v2, minimap2 (v2.28-r1209, (6))). Flye (v2.9.6) (7) was used for assembly with the meta and nano-hq switches, and Polypolish (8) was used to correct assembly errors using illumina data trimmed with AdapterRemoval(v 2.3.4, minlength 100, trimns, trimqualities) (9). BUSCO (v.6.0.0) (10) was used to analyze completeness and annotate core *Plasmodium* genes (plasmodium_odb12, 4203 orthologs). In order to evaluate the number of genomic positions with polymorphisms in the base data, *Plasmodium*-filtered illumina reads were aligned to the polished assembly using BWA (11) and reads mapping within the annotated core genes were filtered using SAMtools and BEDtools (12).

*Processing of existing datasets of ape P. vivax-like samples*

Since only two ape *P. vivax*-like assemblies currently exist at the NCBI genomes database^[1]^, and in order to populate the ape *P. vivax*-like tree, raw illumina data from several studies was downloaded from SRA (see Supplementary Table 1), and KrakenUniq (3) was used to filter out polymicrobial infections by accepting only reads with specific *P. vivax* or ape *P. vivax*-like kmers, as well as all children and parent reads (see Supplementary Table 3 for a list of the used taxIDs). The extracted reads were rarefied to 100x coverage using RASUSA (v.3.0.0) (13) and assembled using SPAdes (v.4.2.0) (14) with the single-cell switch to allow for the expectedly differential coverage resulting from a range of amplification schemes used. The completeness,contamination and annotation of the assemblies was evaluated using BUSCO (v.6.0.0). Getphylo (v.1.0.2)(15) was used to generate a multiprotein phylogenetic tree with the reference genome SalvadorI (GCF_000002415.2) used as seed, a random seed number of 111, a subject and query coverage of 90% of the length, a maximum length of 6000AA, a minimum length of 125 AA, and a minimum presence of a protein in 90% of input sequences, and the build-all parameter to build individual protein trees of all orthologues which satisfy the parameters. In addition to the PvSY43 and PvSY56 which are correctly deposited at NCBIs genbank, the Pvl01 assembly also exist at the NCBI genomes database, but under the wrong species ID “*Plasmodium cf. malariae*” why we did not identify or use the genbank version for this study, but it is identical to the Pvl01 used. NCBI has been contacted to rectify the wrong species assignment. Both the Pvl01 and Pvl06 assembly was downloaded from Figshare and included in the analysis.

Getphylo was run using DIAMOND (v.2.1.13)(16), MUSCLE (v.3.8.1551)(17) and FastTree (v.2.2.0)(18) to identify single copy orthologues, to produce protein sequence alignments, and to generate approximately-maximum-likelihood phylogenetic trees using the default FastTree JTT+CAT models. FastTree calculates the local support values with the Shimodaira-Hasegawa test that are used in Figure 1 as SH-like local supports.

In Supplementary Table 1 column N and O, the number and percent of proteins for each sample used for the tree in Figure 1 is indicated, showing 97.75% presence in the Pvl-KarSSI sample, and generally few samples below 90%, including samples Pvl06 and SYpte90 at 60.11% and 60.96%, respectively. No grouping of the low presence samples is observed.

The concatenated protein tree from getphylo was used for Figure 1. Proteins identified by getphylo were then sorted by the number of varying positions using the AMAS (19) summary function, and the two individual protein alignments with the most sites of variation was used for the example phylogenetic trees in Supplementary Figure A. All phylogenetic trees were visualized using ITOL (20) with nwk tree file outputs of getphylo (Figure 1, Supplementary Figure A). A reduced tree was constructed with getphylo using only samples with BUSCO scores over 60% complete single copy, and the best ape *P. vivax*-like group 2 (21) sequence (p1521_PL_Gabon, SAMN18652601, single copy BUSCO score of 54.9%), and 100% presence (Supplementary Figure B).

After appending BUSCO descriptions to the ID field, BUSCO results gffs were used to annotate the assembly fastas using CLCgenomics (v.25.0.3). Annotated assemblies can be found in Supplementary Dataset I.

*Traveler CAR comparison*

To compare Pvl-KarSSI with the existing resources of the first report of a human infection with ape *P. vivax*-like (TravelerCAR) (22), we mapped all Pvl-KarSSI illumina reads to a database of human (GRCh38.p14) and Traveler CAR sequences (JX444721, partial mitochondrion; KC750243, blood stage SSU; KC750248, CSP) using BWA. We then filtered out secondary hits and human hits using SAMtools, and visualized the mapping on Traveler CAR sequences in CLCgenomics, which was also used for generating consensus sequences.

*Reticulocyte binding protein*

We identified *rbp* genes in Pvl-KarSSI using stand-alone BLASTN searching a database comprised of Pvl-KarSSI contig sequences. We found the Pvl01, PVSY43 and PvSY56 sequences using BLASTN at NCBI to search Whole-genome shotgun contigs (WGS) of *Plasmodium* species (taxid 5820). We found the PvT01, PvC01 and PvP01 sequences using stand-alone BLASTN searching a database comprised of their assembled chromosome sequences. The sequences were aligned using CLUSTALW.

Detailed results and discussion

*Read and assembly statistics*

A total of 45,727,317 Illumina Nextseq1000 read pairs after trimming were generated, of which 6,689,263 read pairs were classified as *Plasmodium* or unclassified and used for polishing and polymorphism evaluation. An Illumina assembly was also attempted, but abandoned as a minimum of 6000 contigs were generated, but the BUSCO score was as high or higher than that of the long reads first approach.

After filtering out human and non-*P.* *vivax* mapping reads, 359,459 Oxford Nanopore 10.4 reads with an N50 of 4,822 nt were used for flye assembly, allowing for the expected differences in coverage depth using the meta switch. After manual removal of Oxford Nanopore adapters and barcodes in the assembly, and after removal of remnants of contaminants by the NCBI submission filter, the assembly consisted of 21,142,361 nt distributed across 473 contigs with an N50 of 110,576 nt.

To evaluate polymorphisms in the dataset, we focused on the annotated single copy core genes from BUSCO, and counted non-majority alternate bases only in those. In order to avoid sequencing errors affecting analysis, only sites with at least 5 identical ALT SNPs were counted. In the 10,054,688 nt single copy core genes including introns, 9,515,910 nt have a coverage of at least 10. Of those 9.5 Mnt, 420 nt has ALT SNPs between 0.1% and 50% of REF, or 0.0044% of positions (see Supplementary Dataset II). We consider this a very low degree of polymorphism, indicating that the patient did not have mixed infections which could impair phylogenetic placement.

BUSCO core protein analysis show that Pvl-KarSSI has 90.5% complete single copy BUSCO orthologs (3802/4203), 2.2% complete duplicated (93/4203), 2.0% fragmented (86/4203), and 5.3% missing (222/4203). This is similar to the best other available ape *P. vivax*-like genomes (PvSY43, , PvSY56 and Pvl01, see Supplementary Table 1).

*Existing datasets of ape P. vivax-like samples*

After initially not realizing that mixed infections existing in many ape samples, KrakenUniq filtering to only keep *P. vivax* and ape *P. vivax-*like and their parents and children TAXIDs reduced the number of duplicate BUSCO models to a maximum of 1.1% of BUSCOs, from 20-40% duplicates before *P. vivax*-filtering in mixed-infection samples (data not shown). This indicates successful filtering for dual infections with other *Plasmodium* sp. Simultaneously, only single-copy complete BUSCOs were used for analysis, further weeding out potential dual infection artefacts. See BUSCO scores for all assemblies in Supplementary Table 1. The getphylo presence/absence table and other results can be found in Supplementary Dataset III.

*Example phylogenetic trees*

The patient had most recently visited Indonesia, where they could have been exposed to both human *P. vivax* and a zoonotic macaque parasite, such as *P. cynomolgi or P. knowlesi*. To evaluate whether the position of Pvl-KarSSI in the tree estimated from concatenated protein sequences could reflect an artefactual combination of sequences from two different species (rather than ape *P. vivax*-like), we inspected two individual protein trees which were used as part of Figure 1. In Supplementary Figure A, subplot 1, a phylogenetic tree built from the getphylo alignment of a conserved *Plasmodium* protein with unknown function is shown (BUSCO:24407at5820). This protein has 151 sites of variation, more than any other individual protein of the 356 proteins used for Figure 1 (see Supplementary Table 4 for the names and descriptions of the 356 BUSCOs). In subplot 2, the getphylo phylogenetic tree of the putative cysteine protease ATG4 is shown, with 133 sites of variation in the amino acid alignment (BUSCO: 24410at5820). This is the protein with the second most sites of variation of the 356 used for Figure 1 (Supplementary Dataset III). Both example phylogenetic trees show all human *P. vivax* sequences grouping in a single clade with support of 1, and the Pvl-KarSSI sample grouping with ape *P. vivax* samples. If the position of Pvl-KarSSI in Figure 1 was a result of a mixed infection of a human *P. vivax* and a macaque parasite, we would expect any individual protein from Pvl-KarSSI to either group with human *P. vivax* or be placed well outside the *P. vivax* clade entirely. Combined with the low BUSCO duplicate score and the low polymorphism count, we consider this evidence that Pvl-KarSSI belongs to the ape *P. vivax*-like groups.

*Reduced tree*

To explore whether the branching order in the phylogenetic tree in Figure 1 was influenced by the parameters used, we generated a reduced phylogenetic tree, only using samples with >60% complete single copy core BUSCOs as well as the best group 2 sequence (54.9% complete) and a presence of 100%. The reduced tree was constructed from 558 loci (compared to 356 for Figure 1), but included only seven ape *P. vivax*-like strains in addition to PvlKarSSI; this tree has identical main clades to the complete tree, with support of 1 for all main clades, and again placing Pvl-KarSSI in ape *P. vivax*-like group 2 (Supplementary Figure B).

*Comparison between TravelerCAR and Pvl-KarSSI*

By mapping all Illumina reads to the limited data from the previously reported case of human infection with ape *P. vivax*-like, we were able to extract a near-complete mitochondrial sequence (5844 nt) and a partial 18S blood stage SSU rDNA sequence. In contrast, only small parts of the CSP gene have coverage (Supplementary Figure C, Supplementary Dataset IV). The number of positions of variation is not sufficient for solid phylogenetic placement within the *P. vivax* and *P. vivax*-like diversity using these sequences, but we did observe several differences between the TravelerCAR and Pvl-KarSSI samples.

*Reticulocyte-binding protein analysis*

Loy et al. (23) found that three of the genes (*rbp2d*, *rbp2e* and *rbp3)* in the reticulocyte-binding protein gene family are pseudogenes in all strains of human *P. vivax*, but are conserved as intact, presumably functional, genes in strains of *P. vivax*-like from apes. Thus, analysis of the *rbp* genes provides a complementary approach to distinguish between strains of *P. vivax* from humans and *P. vivax*-like from apes. Here we compare sequences from Pvl-KarSSI to the three assembled ape *P. vivax*-like genomes and to three representative *P. vivax* genomes from humans:

PvSY43:	*P. vivax*-like from a chimpanzee (Loy et al. 2018) (23)

PvSY56:	*P. vivax*-like from a chimpanzee (Loy et al. 2018) (23)

Pvl01:	*P. vivax*-like from a chimpanzee (Gilabert et al. 2018) (24)

PvT01:	*P. vivax* from a human in Thailand (Auburn et al. 2016) (25)

PvC01:	*P. vivax* from a human in China (Auburn et al. 2016) (25)

PvP01:	*P. vivax* from a human in Indonesia (Auburn et al. 2016) (25)

*rbp3*

The *rbp3* gene is on chromosome 14 in human *P. vivax*, and was found on contig_40 in our genome assembly. The three assembled human *P. vivax* genomes (strains P01, T01 and C01) have two premature stop codons, similar to 82.4% of 227 strains examined previously (23).

The three assembled *P. vivax-*like genomes, from strains SY43, SY56, and Pvl01 (from PmlGA01), have Arg (AGA) and Ser (TCG) codons at the sites of the stop codons (TGA, TAG) in human *P. vivax.* The *rbp3* gene sequence in contig_40 has the same Arg and Ser codons as the other *P. vivax*-like strains.

*rbp2e*

The *rbp2e* gene is on chromosome 7 in human *P. vivax*, and was identified on contig_50 in the Pvl-KarSSI assembly.

In *P. vivax*-like strains SY43, SY56, and Pvl01, the gene comprises a first exon of 55 nt, an intron of 288 nt, and a second exon of 8144 nt. In human *P. vivax* strains P01 and T01 there are four indels, of either 1 nt or 4 nt, which all give rise to frameshifts. These are an insertion (4 nt) near codon 156, a deletion (4 nt) near codon 285, a deletion (1 nt) near codon 1878, and an insertion (1 nt) near codon 2273. All four indels were previously found in 28.4% of 162 strains of human *P. vivax* (23). Strain C01 lacks the deletion near codon 285; similarly, only three indels were found in 21.3% of 162 strains of human *P. vivax* (23). None of these four indels are present in *P. vivax*-like strains SY43, SY56, and Pvl01. Also, codon 881 in the three *P. vivax*-like strains (SY43, SY56, Pvl01) is GAA (encoding Glu), but in the three human *P. vivax* strains (P01, T01, C01) there is a mutation to TAA (a stop codon); as previously found in all strains (23).

In contig_50 in the Pvl-KarSSI genome assembly, only the last 7347 nt of exon 2 are found, at the beginning of the contig, meaning that the first 284 codons are not included in the contig. Thus, we cannot comment on the site of the human *P. vivax* indel near codon 156. However, it is clear that the deletion near codon 285 is not present in contig_50, and the two other human *P. vivax* indels are not present either. Also, the site homologous to codon 881 is GAA in contig_50 in our genome assembly.

*rbp2d*

The *rbp2d* gene is near the end of chromosome 14 in *P. vivax*. The reference strain *P. vivax* P01 has 727 protein-coding genes annotated on chromosome 14; *rbp2d* is gene 711. The orthologue of gene 692 is at the end of contig 40, but there is no homology to genes 693-727 in the contigs, indicating that the region encompassing the *rbp2d* gene is missing from the Pvl-KarSSI genome assembly; this is likely because of a lack of coverage in the Oxford Nanopore data, which is consistent with the observation that chromosome ends are poorly captured in our dataset.

**Supplementary Figures legend:**

Supplementary Figure A: Two example phylogenetic trees of individual protein alignments of the 356 used for Figure 1. Subplot 1: Protein with unknown function (BUSCO:24407at5820), Subplot 2: Putative Cysteine protease ATG4 (BUSCO: 24410at5820). In both examples, all human *P. vivax* strains form a clade with support of 1, and the Pvl-KarSSI is placed within the ape *P. vivax*-like clades. This is evidence that Pvl-KarSSI is not a result of a mixed infection.

Supplementary Figure B: Phylogenetic tree built from a reduced dataset, using only HQ samples as well as the best ape *P. vivax*-like sequence from group 2. Only proteins identified in all samples (n=558 BUSCO proteins) were used for the tree. The main clades are identical to the ones in the full tree in Figure 1 of the main text, and have a support of or near 1, indicating that adding proteins or removing samples do not affect the main structure or interpretation of the Figure 1 tree.

Supplementary Figure C: comparison between the previous human case of ape *P. vivax*-like “TravelerCAR” and the Pvl-KarSSI strain presented here. Subplot 1: mapping of Pvl-KarSSI reads on the TravelerCAR reference blood stage 18S SSU. Much of the 18S ribosomal fragment from the TravelerCAR sample is covered. Multiple SNPs between the samples are observed, but the cause (technical or biological) and implications of them remain unclear. Subplot 2: mapping of Pvl-KarSSI reads on the TravelerCAR mitochondrial sequence. Subplot 3: the CSP gene was not found to have good coverage in the Pvl-KarSSI sample, but several SNP differences can be observed in the brief windows with overlap. Subplot 4: section of alignment of Pvl-KarSSI, TravelerCAR and multiple other partial 18S SSU sequences from the blood stage. While too few differences exist for solid analysis, the window shown (100/~1000 nt) highlights a locus where the two samples Pvl-KarSSI and TravelerCAR differ, and each profile is seen in other samples.
