## Supplementary Figure A for "Human infection with an ape *Plasmodium vivax*-like parasite"

Subplot 1 Conserved *Plasmodium* protein of unknown function (BUSCO:24407at5820)

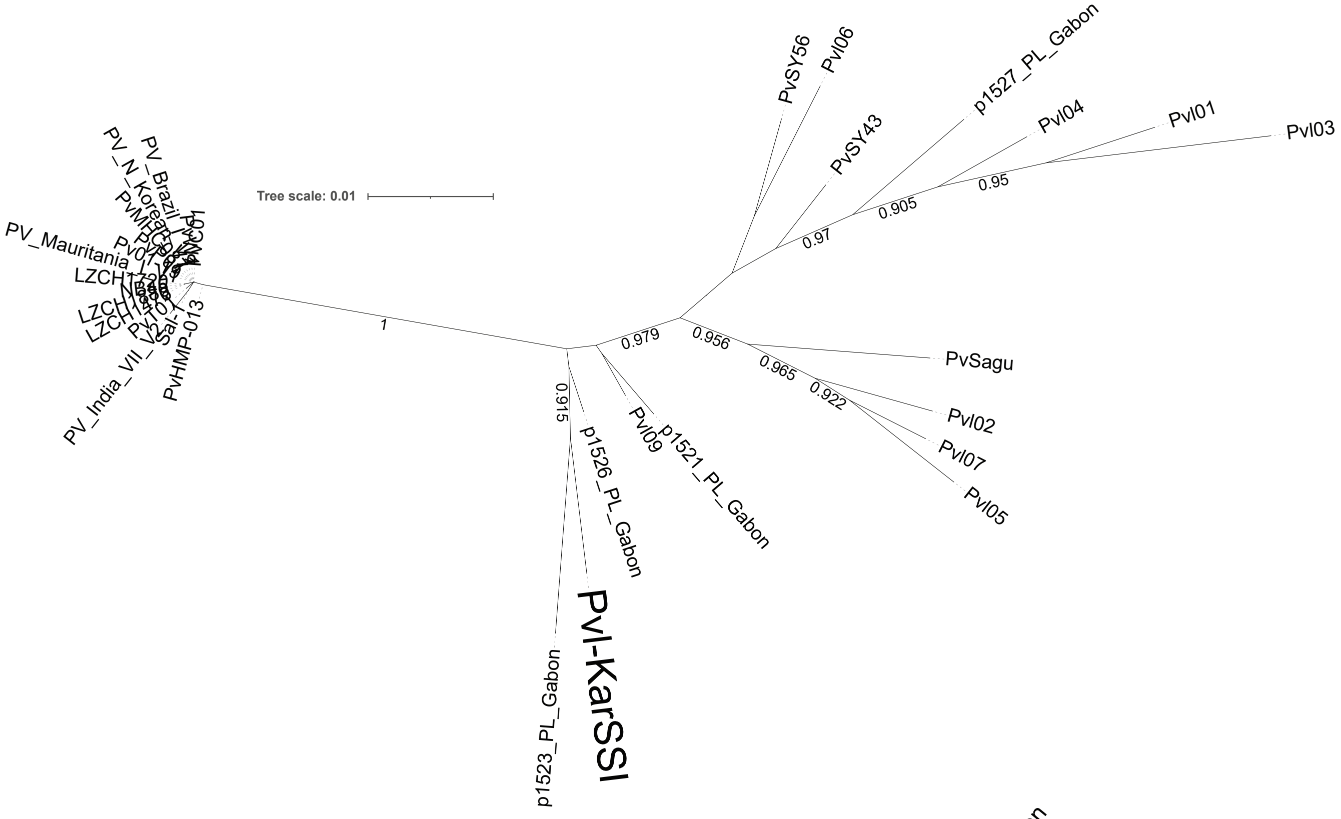


Subplot 2 Putative cysteine protease ATG4 (BUSCO:24410at5820)

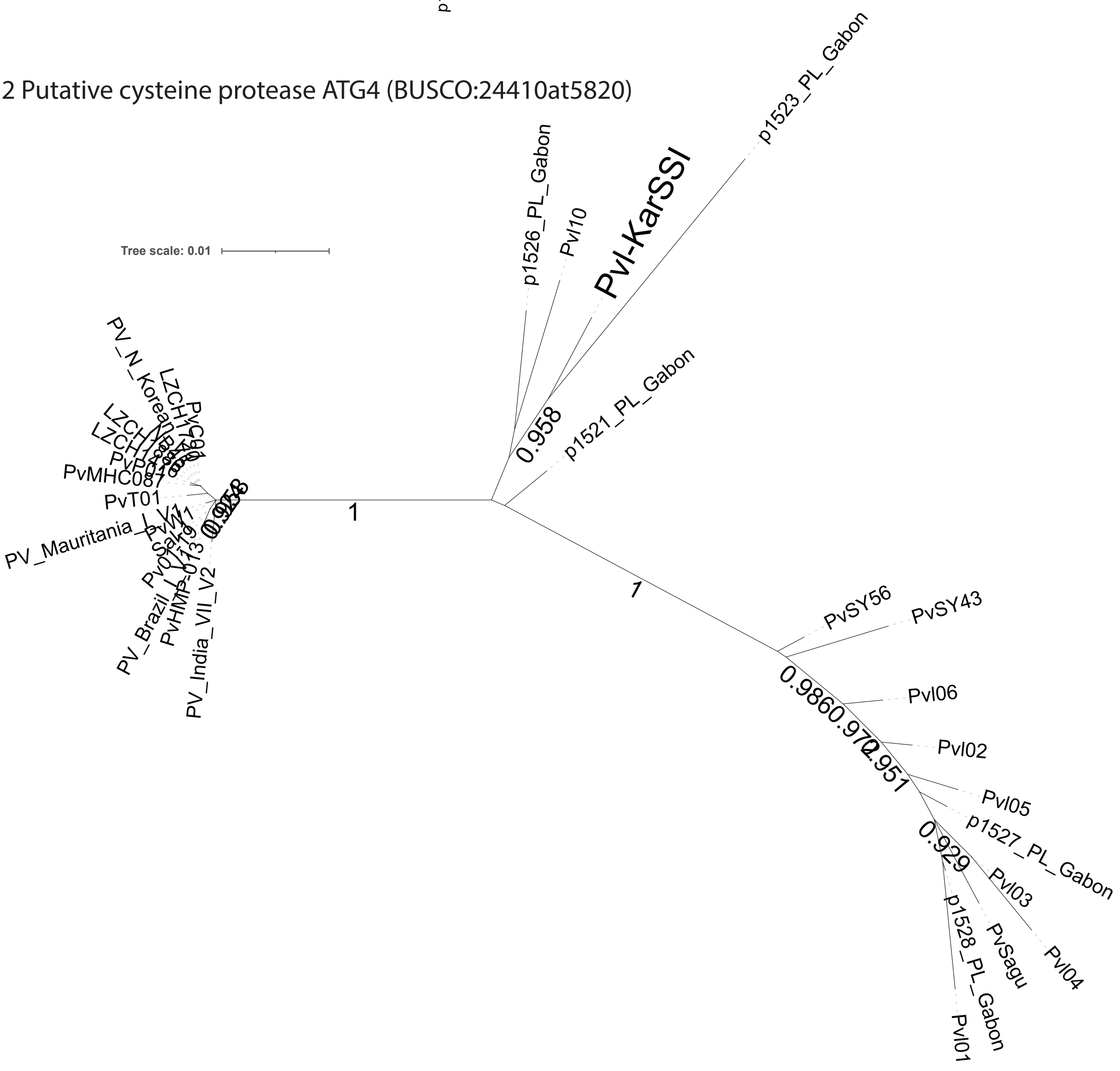
