## Supplementary Figure C for "Human infection with an ape *Plasmodium vivax*-like parasite"

### Subplot 1 blood stage 18S SSU comparison of PvI-KarSSI reads and TravelerCAR

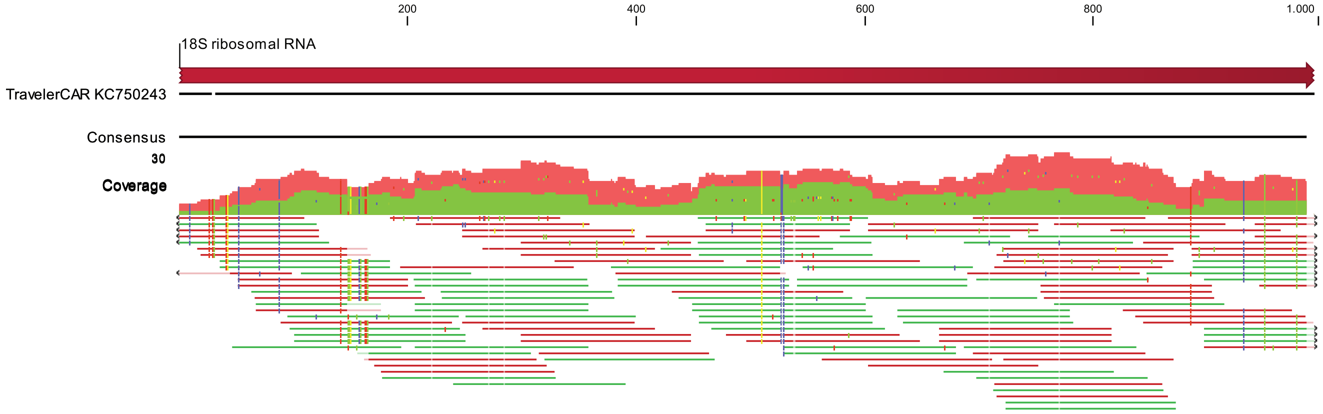


### Subplot 2 mitochondrion comparison of PvI-KarSSI reads and TravelerCAR

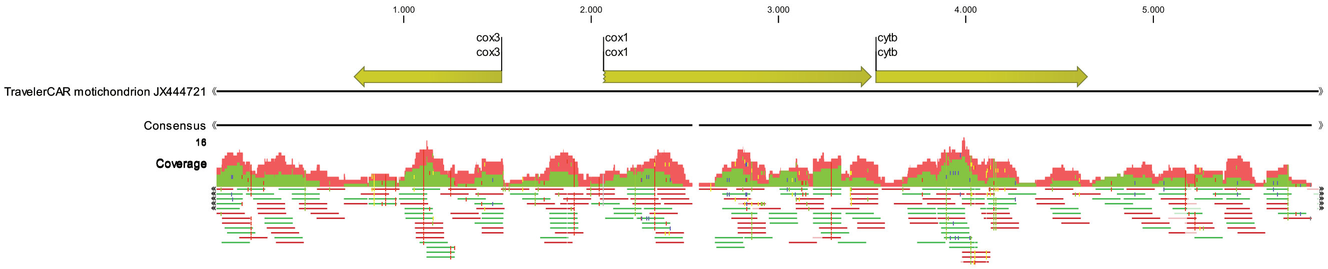


### Subplot 3 CSP comparison of PvI-KarSSI reads and TravelerCAR

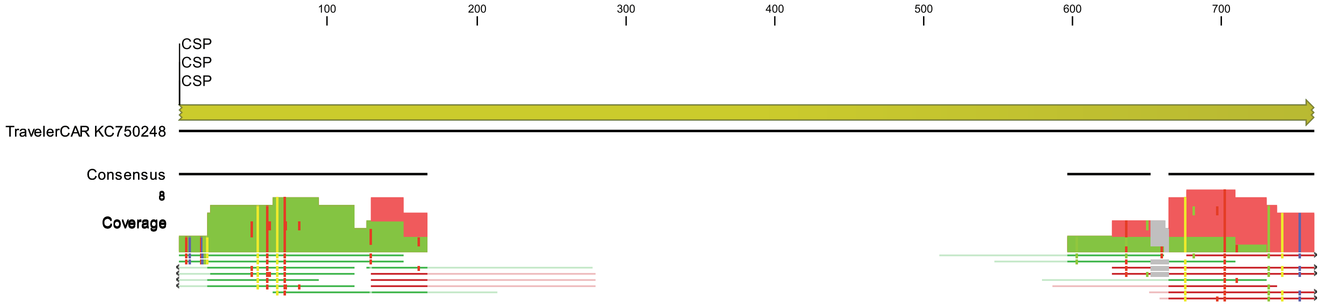


### Subplot 4 section of 18S SSU alignment focusing on incongruence

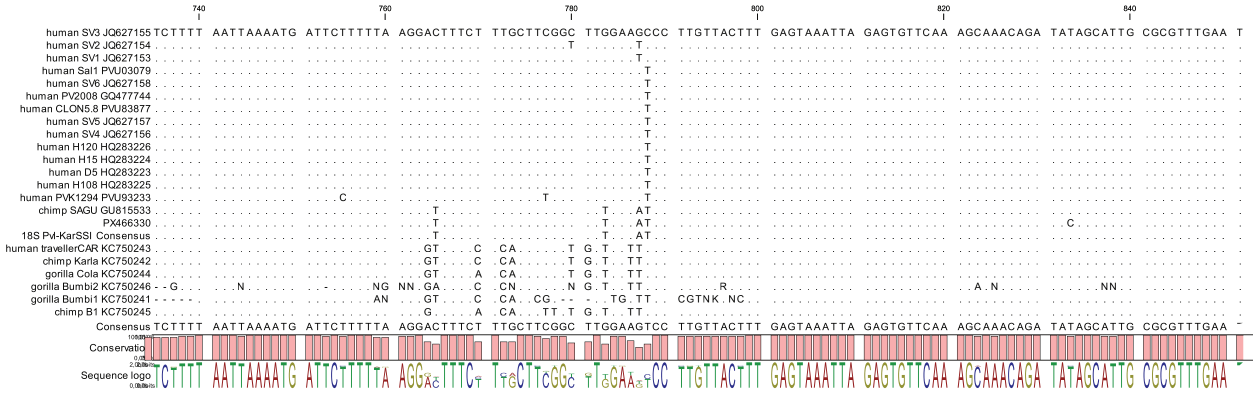
