## Supplementary figures and images for "Human infection with an ape *Plasmodium vivax*-like parasite"

### Supplementary Figure B

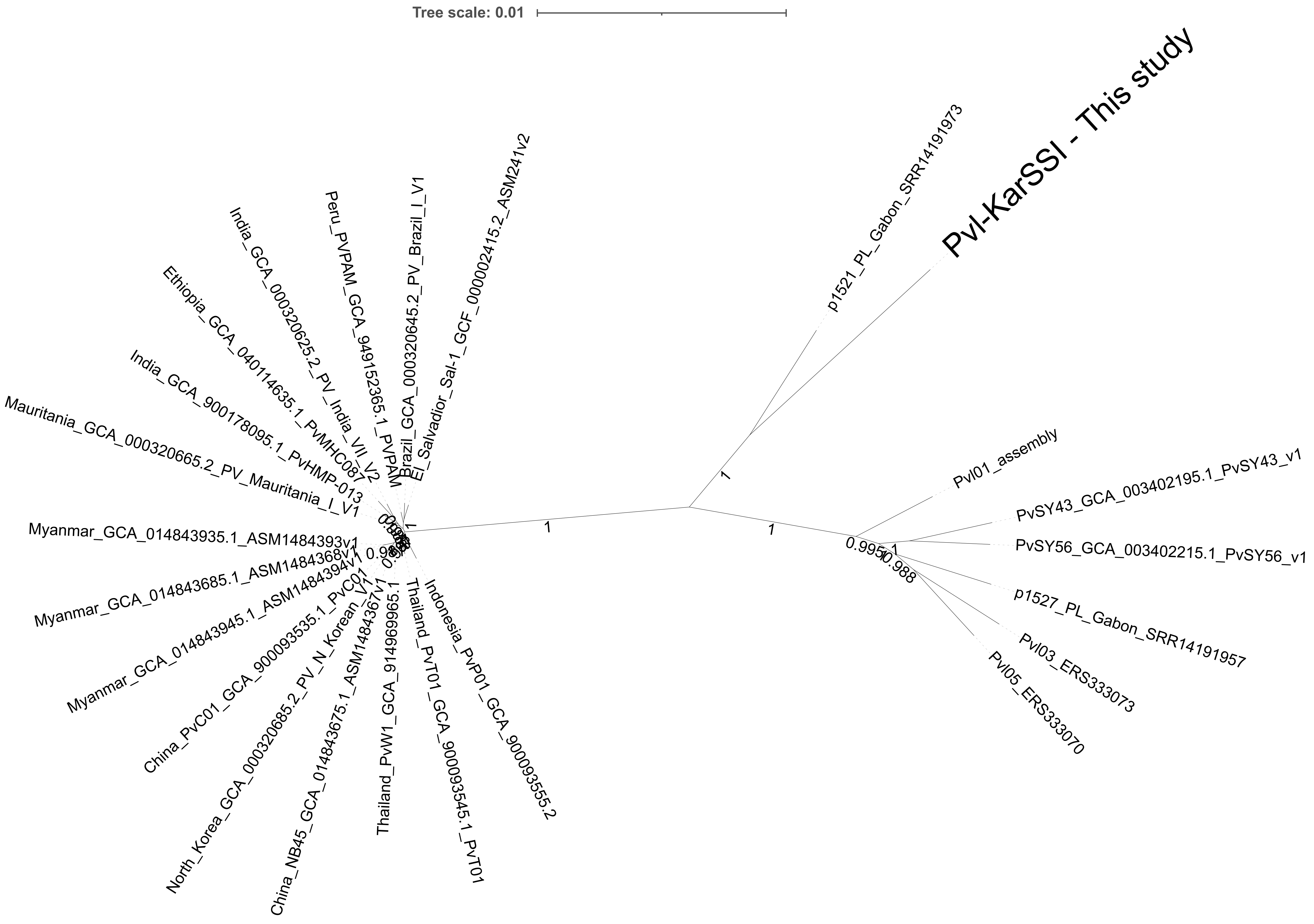
